# Three-dimensional tissue platform co-laid with native collagen fibers and cells for phenotypic screening of stem cell interactions

**DOI:** 10.1101/2025.02.28.640824

**Authors:** Rui Tang, Aixiang Ding, Chen Fu, Kentaro Umerori, Marvin Rivera, Daniel S. Alt, Christopher M. Carmean, Li Li, Steven J. Eppell, Anthony Wynshaw-Boris, Eben Alsberg

**Author notes:** (RT), (EA).

## Abstract

Phenotypic screening of cell-cell and cell-matrix interactions is critical yet challenging for drug discovery and disease modeling. In this study, a scalable 3D tissue platform was developed by co-laying extracted natural insoluble collagen fibers, mesenchymal stem cells, endothelial cells, and neural progenitor cells for phenotypic screening. Cell growth and interactions were enhanced in the co-laid platform, evident through increased cell proliferation, viability, and vascularization. Dense vascular networks rapidly formed through cell-cell and cell-matrix interactions without adding a traditionally needed growth factor set. Both *in vitro* and implantation studies confirmed that these blood vessels were of human origin. To evaluate the phenotypic screening of cell-cell and cell-matrix interactions, we propose a phenotype screening prototype for stem cell interactions that utilized multivariate analysis encompassing both cell-cell and cell-matrix interactions and demonstrated its effectiveness to screen vasculature formation and autism spectrum disorder (ASD) models. Using the prototype, we confirmed that collagen crosslinking, ROCK, WNT, and YAP pathways impact vasculogenesis. In addition, ASD donor-derived neural progenitor cells can be distinguished from non-ASD control donor-derived neural progenitor cells.

## Introduction

Phenotypic screening, a system focusing on the effects of biological systems, is a classical method that once dominated drug discovery in the past. After being supplanted by target-based drug discovery for decades, recent technological advances that capture phenotypic changes in cells and tissues previously unquantifiable or overlooked have enabled a renaissance of phenotypic screening, resulting in multiple drug approvals in recent years.^1^ Despite promising expansions in the capabilities of these systems, developing workflows for screening and processing hits that are both efficient and effective remains a major challenge.^2^ Platforms that can enrich the phenotypic changes and transduce the signals for detection and analysis still need further development. Ideally, multiple variables need to be extracted from the screening outcomes. These variables can then be analyzed separately or holistically to characterize the phenotypic differences of cells or tissues under different conditions.

Tissues engineered from stem cells in three-dimensional (3D) better recapitulate natural tissue growing conditions compared to two-dimensional (2D) culture^3^ and, therefore, hold great promise for phenotypic screening for disease modeling and drug discovery.^4^ In an optimized 3D tissue environment, phenotypic changes are often stronger compared to 2D counterparts due to stronger cell-cell and cell-extracellular matrix (ECM) interactions and thus can be captured and analyzed more easily. Various 3D culture systems have been developed and employed in tissue engineering, disease modeling, and drug discovery.^5^ While natural growing environments usually possess both strong cell-cell and cell-ECM interactions, most artificial 3D systems often elevate one type of interaction at the expense of another. For example, in hydrogel systems, regardless of the density of embedded cells, individual cells are predominantly surrounded by hydrogel materials rather than other cells. Hydrogels need to be degraded when strong cell-cell interactions are required, such as for neural tissues.^6^ Populating cells into pre-fabricated and natural decellularized scaffolds relies on cell infiltration. Cell density inside these scaffolds depends on scaffold pore sizes and depth,^7^ and the influx of migratory cells is problematic.^8^ On the other hand, it is difficult to evenly introduce ECMs into densely packed cell aggregates. As a result, *in vivo* cell-ECM interactions are challenging to replicate *in vitro* using the organoid technique.^9^

In an ideal *in vitro* 3D tissue culture system, maintaining balanced cell-cell and cell-ECM interactions for individual cells is critical to mimic the *in vivo* microenvironment. In nature, cells usually reside in a hierarchical ECM that is highly ordered.^10^ An abundant ECM family of proteins especially collagens support a variety of cell types in most human organs and tissues.^11^ Type I collagen molecules assemble into fibrils with diameter of a hundred nanometers or so, which are bundled into fibers with diameters of about 10-15 μm in average particularly in dermal tissues.^12,13^ Collagen fibers provide sufficient mechanical, chemical, and biological support to cells. Nevertheless, few scaffolds have been developed to sufficiently imitate the natural collagen fiber for 3D cell growth and tissue engineering. Re-assembly of collagen molecules *in vitro* often results in collagen hydrogels or sponges consisting of fibrils but not fibers. Although such matrices have been widely used and have proven to support cell growth, due to the fine size of the fibrils, cells are usually surrounded and trapped by tiny fibrils.^14^ Reducing the fibril density increases the pore size thus increasing cell-cell interaction but at the cost of reduced mechanical strength of the matrix.

Since collagen fibers are one of the major natural ECM structures to support cell growth, we hypothesize that directly isolating these collagen fibers from tissues for use as tissue engineering matrices potentially provides a microenvironment for resident cells that better mimics natural conditions. In this study, we dissociated a collagen-rich tissue scaffold into independent collagen fibers (Figure 1a). These collagen fibers are flexible and slurry-like yet robust. Traditional water-lay techniques are widely used for paper and non-woven fabric manufacturing, which disperse fibers in water and lay them on the meshes by filtering the water out to form the product, and provide a method with the potential for scalability. Adapting this technique, we dispersed collagen fibers with cells in the culture media, and co-laid them through centrifugation to construct tissues rapidly while maintaining effective cell-cell and cell-ECM interactions. Based on this technique, we propose a 3D tissue-level phenotypic screening platform that efficiently detects the changes in cell-cell and cell-ECM interaction. We also show the uses of this proof-of-concept by phenotypically distinguishing the vasculature formation of endothelial cells under different stresses and autism spectrum disorder (ASD) donor cell behaviors compared to those of non-ASD controls.

## Results

### Collagen fiber preparation and characterization

In the first step, decellularized, purified, and ground calf skin powders (Figure 1b) were chosen to extract the dispersible collagen fibers. The skin powders were dissociated with pepsin under 4 °C, similar to a previous report.^15^ However, instead of fully degrading the tissue scaffold to soluble collagen molecules, the dissociation was controlled to the degree that massive fiber bundles were liberated. Sequentially, collagen fibers were purified by gradual centrifugation. The collagen fibers extracted were 11 ± 6 μm in diameter and hundreds of microns in length, in agreement with the size of those in natural tissues,^13^ down from 126 ± 71 μm in diameter of the skin powders (Figure 1b, c; Supplementary Figure 1a, b). The fibers contain about 76% Type I collagen, 16% Type III collagen, and 8% Type II keratin, according to proteomic analysis (Supplementary Table 1). The collagen fiber structure was further assessed by atomic force microscopy (AFM). Low magnification AFM revealed that the micron diameter fibrous structures consisted of bundled sub-micron fibrils (Figure 1d). High magnification AFM confirmed intact D-banding of the collagen fibrils (Figure 1e; Supplementary Figure 1). Notably, the collagen fibers form slurries in aqueous solutions that can be easily expelled through a 16-gauge needle by a syringe (Figure 1f, Supplementary Video S1). This feature enables rapid preparation of tissues in multi-well plates and for large-scale use.

**Figure 1.**
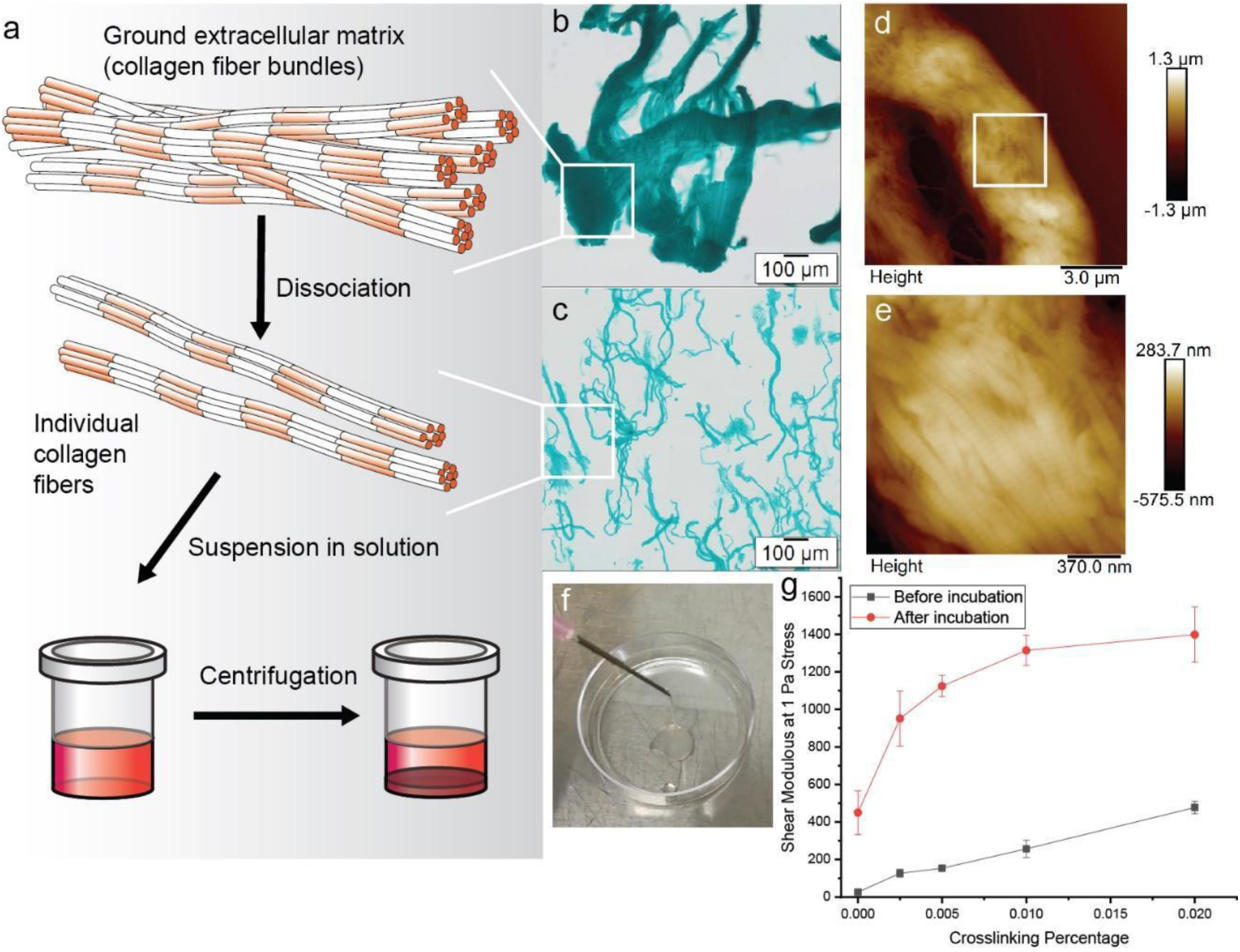
Preparation and characterization of extracted natural insoluble collagen fibers. a, Scheme of the preparation process. b, The ground calf skin is stained by Fast Green. c, Isolated collagen fibers stained by Fast green. d, AFM of an individual collagen fiber showing the fibril bundle structure. e, AFM of the collagen fiber at high magnification showing D-bands. f, Extrusion of collagen fibers through a 16-gauge needle. g, Rheometry results of uncrosslinked and crosslinked collagen fibers with or without incubation at 37 °C for 24 hr.

Crosslinking of collagen alters its mechanical and chemical properties, thus providing a tunable microenvironment for cell growth. However, after crosslinking, traditional collagen materials can no longer be readily suspended in aqueous solution. Instead, gels or solid scaffolds are the usual products. One advantage of collagen fibers is that they can be facilely pre-crosslinked and remain suspended until tissue construction. In the current study, collagen fibers were pre-crosslinked by glutaraldehyde at different weight percentages, from 0 to 25%. The rheometry showed that with the increase of the crosslinker concentration, the shear moduli of the materials gradually increased (Figure 1g). Incubation of the collagen fibers at 37 °C for 24 hr further increased the shear moduli of the uncrosslinked and crosslinked collagen fibers. Since the shear modulus describes the rigidity of material under shear stress, the rheometric results confirmed that the stiffness of the collagen fibers increased with crosslinking.

### Tissue construction using collagen fibers

To verify the biocompatibility of collagen fibers, two cell lines, NIH/3T3 and HeLa, were co-laid with the collagen fibers through mixing and centrifugation-based laying in media to form a tissue (Figure 2a). The tissues were then cultured for seven days, and cell viability was tracked through laser scanning confocal microscopy (LSCM) and DNA assay. After 24 hr, both cell types were well distributed in the formed tissue (Figure 2 b and c). To further confirm that the cells resided in the collagen fiber, one tissue containing NIH/3T3 cells was torn and imaged after 48 hr of culture (Figure 2d). From the brightfield image, the NIH/3T3 cells adhered to the collagen fibers, providing evidence of the successful residency. LSCM images (Supplementary Figures 2 and 3) and DNA quantification (Figure 2e) also indicated that both cell types proliferated during the 7-day culture. In contrast, the ground calf skin, without dissociation, did not harbor cells after 24 hr (Supplementary Figure 4), indicative of the necessity to dissociate the tissue to individual collagen fibers for cell lodging. Human mesenchymal stem cells (hMSCs), a type of multipotent stem cells, were co-laid with collagen fibers to form tissue as well. Cells remained healthy for up to 14 days in proliferation media, as revealed by LSCM (Supplementary Figure 5). As demonstrated in Supplementary Video S2, the tissue containing hMSCs can be pulled up by tweezers and transferred to phosphate-buffered saline for further manipulation with preserved geometry, suggesting that the tissue is compliant but strong enough to withstand manipulation. These combined results demonstrate that the processed collagen fiber matrix is capable of supporting a variety of cell types, while the original undissociated calk skin fiber bundles (skin powder) are not.

**Figure 2.**
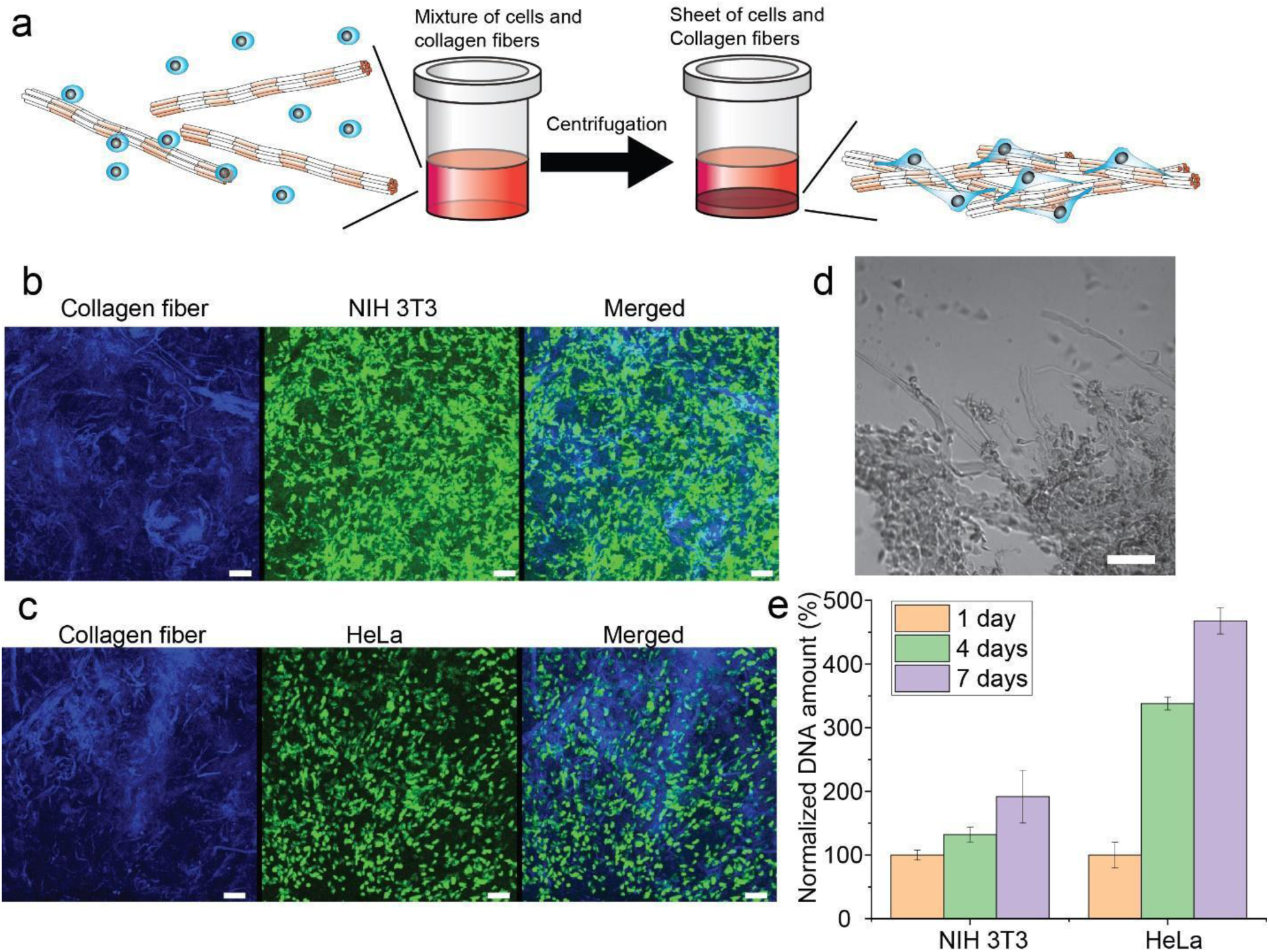
Collagen fibers support the proliferation of cells in the 3D constructs. a, Scheme of 3D construct preparation. b, Z-stack LSCM image of NIH/3T3 cells in the construct after 24-hour culture. Blue: autofluorescence of collagen fibers; green: live cells stained by fluorescein diacetate. Bar: 100 μm. c, Z-stack LSCM image of HeLa cells in the construct after 24-hour culture. Blue: autofluorescence of collagen fibers; green: live cells stained by fluorescein diacetate. Bar: 100 μm. d, Torn region of the construct containing NIH/3T3 cells after 48-hour culture, showing the presence of cells along with collagen fibers. Bar: 100 μm. e, Normalized DNA content assays for NIH/3T3 and HeLa cells growing in the collagen fiber 3D constructs for 7 days. DNA contents after one day were set to 100%.

### Vasculogenesis of the constructed tissues

Since the collagen fibers enable both cell growth, migration, and interactions, this platform is expected to support multiple types of cell interactions needed for complex tissue construction. To test this hypothesis, we co-cultured hMSCs and human umbilical vessel endothelial cells (HUVECs) in the collagen fiber platform for vasculogenesis (Figure 3). hMSCs contain important pericytes,^16^ and they help endothelial cells survive and form vasculatures through the paracrine pathway.^17^ To maximize the influence of the hMSC-HUVEC interaction, the cells were cultured in DMEM supplemented with 10% FBS and 10 ng/mL FGF-2, which is insufficient for HUVECs alone to survive (Figure 3a). However, when cultured in the collagen fiber platform, HUVECs survived and formed vasculature within 3 days (Figure 3a). After a 7-day culture, the vasculature density greatly increased in 3D as evidenced by staining of the endothelial marker hCD31 (Figure 3a, Video S3). To verify the necessity of cell-cell contact, we placed hMSCs on the top layer of a transwell plate and HUVECs on the bottom layer. After 3 days, no HUVECs survived (Figure 3a). This result confirmed that direct contact with hMSCs is vital to maintain HUVEC viability without the addition of extra growth factors presented in standard endothelial expansion media such as Vascular Endothelial Growth Factor (VEGF), Epidermal Growth Factor (EGF), Insulin-like Growth Factor 1 (IGF-1). Co-cultures of these cells in collagen gel or GelMA for 7 days under the same culture conditions did not generate obvious vasculatures (Figure 3a). This result is consistent with the hydrogels wrapping the cells resulting in sterically hindered cell migrations^18^ that disallowed sufficient cell-cell interactions. H&E staining of the sectioned tissue confirmed vascular lumen formation after 7 days (Figure 3b, Supplementary Figure 6). These lumens stained positive of hCD31 and laminin, indicative of *in vitro* blood vessel formation (Figure 3b, Supplementary Figure 6). This result proved that direct contact with hMSCs is vital for the survival of HUVECs and vasculature formation. These results demonstrated that without the supplement of common growth factors for vasculogenesis, such as vascular endothelial growth factor, direct hMSC-HUVEC contact or paracrine signaling must take place for HUVEC survival and subsequent vasculogenesis. Collectively, the collagen fiber platform supports direct cell-cell interactions, allowing paracrine pathways to be maintained. As vascular formation is critical for tissue modeling and engineering, this ability will allow more complex tissue formation for both *in vitro* and *in vivo* use.

We implanted tissues cultured for 3 days *in vitro* onto chick chorioallantoic membranes (CAMs) and incubated them for another 7 days to further explore the formation of the vasculature (Figure 3c). Histological images showed that the implanted tissue was perfused with red blood cells (Figure 3d). It should be noted that avian red blood cells are nucleated.^19^ Therefore, they exhibited dense bright red cytoplasm and dark purple nuclei with H&E staining. Human CD31 immunostaining indicated that these blood cells were surrounded by human endothelial cells (Figure 3e and Supplementary Figure 7). Human ALU element in-situ hybridization strongly stained the blood vessels in implanted tissue compared to those in the adjacent chicken tissues (Figure 3e and Supplementary Figure 8), further confirming that the cells are of human origin. Taking the *in vitro* and implantation tests together, the vasculature that appeared in the engineered tissues are human blood vessels. Strong cell-cell and cell-ECM interactions are sufficient to stimulate vasculature formation without extra growth factors.

**Figure 3.**
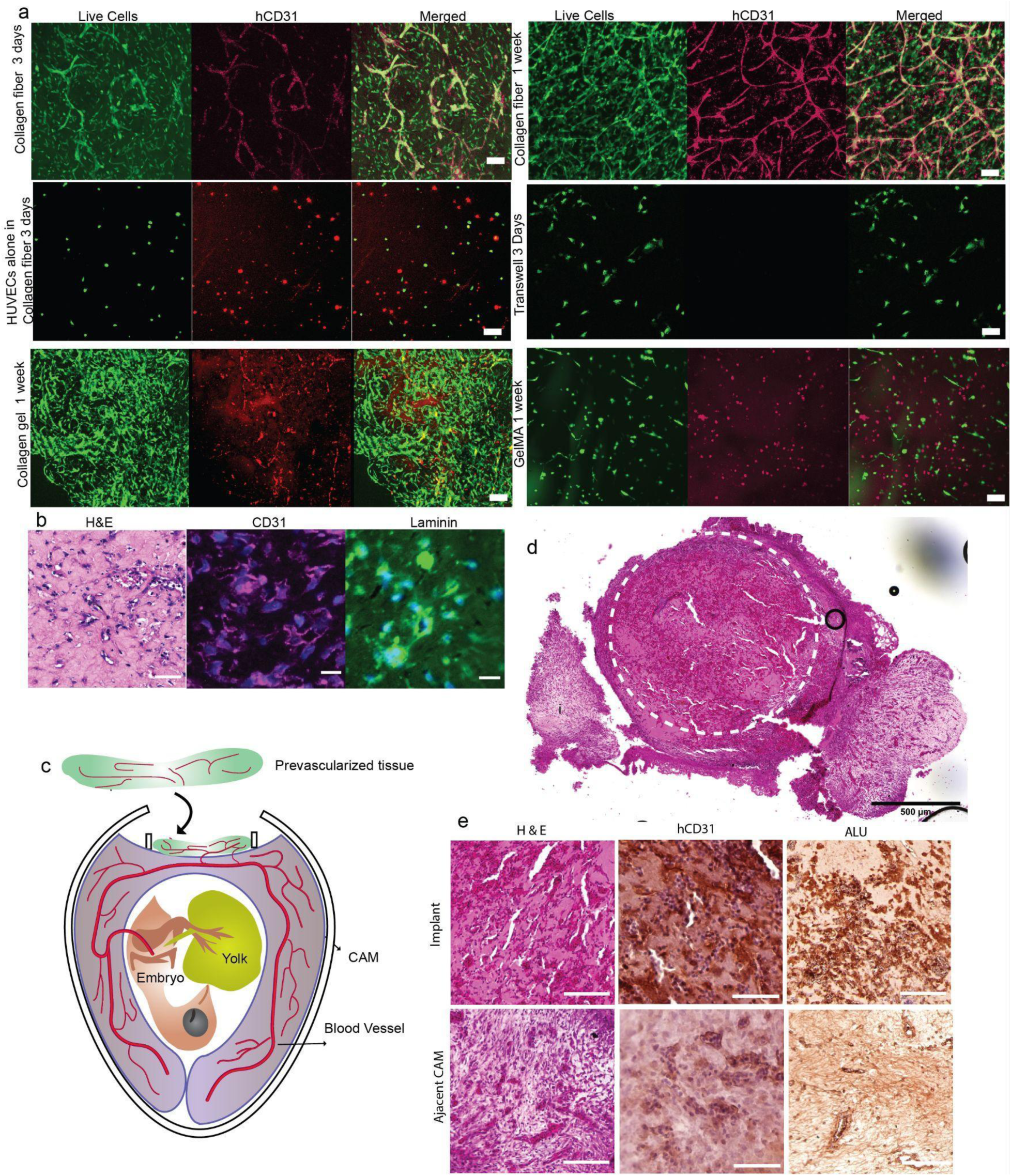
Collagen fibers enable rapid vasculogenesis by endothelial cells through interaction with hMSCs. a, Co-culture of HUVECs and hMSCs in collagen fiber, transwell, collagen gel, or GelMA. For transwell culture, hMSCs were seeded in the transwell insert, and HUVECs were grown on the bottom layer. Bars: 100 μm. b, H&E staining and immunostainings of the sectioned tissue co-cultured for 7 days. c, Scheme showing the transplantation of a vascularized *in vitro* tissue onto CAM for blood vessel connection and blood perfusion. d, large-scale H&E staining of implanted tissue. Bar: 500 μm. e, H&E, hCD31, and ALU staining of regions of interest of implanted tissue. Bars except d: 100 μm.

### Screening platform for factors influencing vasculogenesis

With the demonstration that collagen fiber-based tissues retain strong cell-cell and cell-matrix interactions, we therefore propose a rapid and scalable platform to construct 3D tissues for versatile applications (Figure 4). Within this proposal, a mixture of cells, collagen fibers, and media are transferred to well plates in a batch. The plates are then centrifuged to spin down cells and collagen fibers simultaneously to immediately form the tissue sheets at the bottom of each well. Different cells, collagen fibers with different crosslinking, or drugs of interest can be introduced in the system during or after tissue preparation. During the culture of the tissues, the phenotype characteristics will gradually appear and can be recorded as one or multiple variables. After the collection of a panel of the characteristics, a multivariate analysis can then be performed to distinguish the phenotypes of cells or tissues under certain conditions.

**Figure 4.**
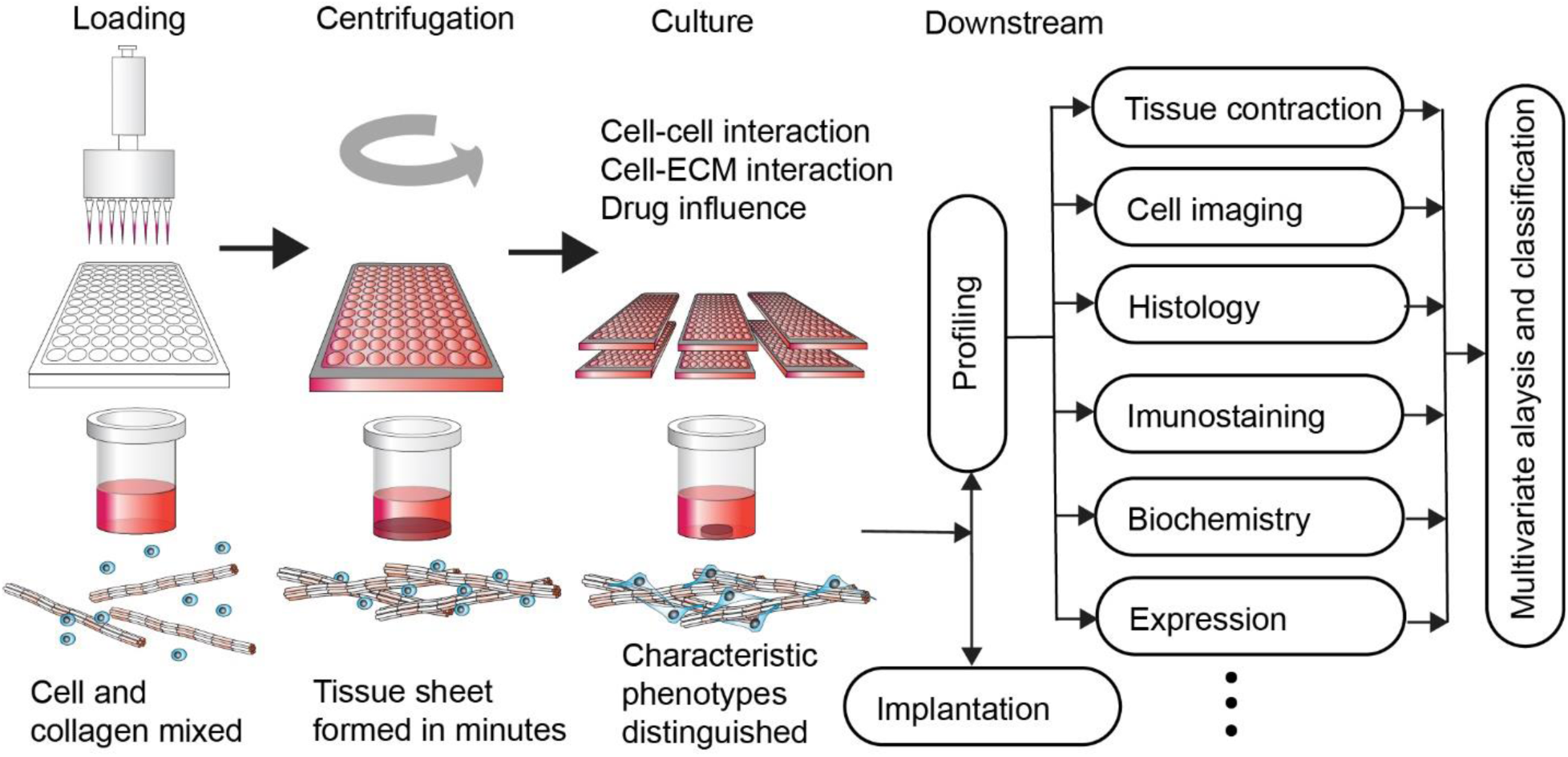
Schematic presentation of the proposed phenotypic screening platform.

We then created a prototype using the hMSC and HUVEC interactions for vasculature formation to demonstrate the concept of the phenotype screening platform (Figure 5). Since cell interactions play important roles in vasculature formation in our tissue system without the addition of multiple endothelial differentiation factors, a reliable phenotypic screening platform should be able to capture the changes quickly if these interactions are altered. Literature has shown that a variety of signals will impact the interactions at different times. Two sets of tests were performed to adjust the interactions in this platform and test their impact. The resulting vasculatures of each individual tissue were stained with anti-hCD31 and imaged using a high-content imager. The images were then preprocessed to highlight the vasculatures and quantitatively analyzed using ImageJ with the vasculature plugin^20^ (Figure 5a). Twenty variables extracted from the analysis were plotted through linear discrimination analysis (LDA), a multivariate analysis method for dimensionality reduction, classification, and extraction of important features of data.

In the first set of tests, collagen fibers were either untreated or pre-crosslinked to different extents to impart different mechanical strengths to the ECM. After 7 days of culture, substantial amounts of vasculature in all groups wase observed through LSCM (Supplementary Figure 9). After 2 weeks of culture, groups with high crosslinking percentages (12.5% and 25%) showed dramatically reduced vasculogenesis compared to other groups through single variable analysis of high-content images (Supplementary Figure 10). Considering all 20 variables together with LDA, the vasculatures of all 5 groups were well distinguished (Figure 5b). This result suggests that the differences in cell-cell interactions induced by cell-ECM interactions can be captured in this phenotypic screening platform.

In the second set of tests, various substances (Figure 5c) were added into the culture on Day 0 or Day 3, impacting the initiation and extension of the vasculature, respectively. The chosen substances include an MMP inhibitor (Marimastat) to minimize ECM modulation of cells, a WNT inhibitor (XAV939) to prevent direct cell-cell interaction, a GSK3 inhibitor/WNT activator (Chir99021) to enhance cell-cell interaction, a YAP inhibitor (Verteporfin) to shelter cells from sensing the mechanical environment, a ROCK inhibitor (Y-27632) to block cell migration, and VEGF to directly inhibit vasculature formation, and VEGF, a known growth factor for vasculature formation. In the presence of the drugs, the morphology and density of the vasculature were visually different (Supplementary Figures 11 and 12). LDA (Figure 5d and f) and pairwise Permutational multivariate analysis of variance (PERMANOVA; Figure 5e and g) clearly highlighted ROCK, WNT, and YAP pathways. Unsurprisingly, the ROCK inhibitor which prevented cell migration significantly decreased the initiation and formation of vasculature, demonstrating that maintaining cell migration is important throughout vasculogenesis. The WNT inhibitor, which downregulates a pathway critical for cell-cell communication, interfered with vasculature initiation but not extension. From this result, we confirmed that the WNT signaling pathway is necessary for initiation of vasculogenesis. Once the initiation is complete, the WNT signaling pathway is suppressed. This result is in agreement with previous findings that WNT signaling is important at early steps of vasculogenesis rather than in healthy and resting vasculature.^21^ The YAP inhibitor downregulated the extension of the vasculature but not its initiation. The YAP signaling pathway is responsible for sensing mechanical cues during vasculogenesis^22^ and reacts to microenvironmental stiffness changes.^23^ This result suggests that mechanical cues played their roles after initial cell interactions occur. It also explains higher stiffness tissues consisting of pre-crosslinked collagen fibers have vasculature patterns distinguishable from their lower stiffness counterparts. The MMP inhibitor had no obvious effect on vasculogenesis of the tissues, suggesting ECM degradation has little impact on vasculogenesis under experimental conditions. The WNT activator and extra VEGF signal also showed no effect on vasculogenesis of the screened tissues, suggesting that these pathways were already turned on. This result is not surprising given their importance during vasculogenesis. The results confirmed that inhibition of WNT, YAP, and ROCK pathways at different times will impact vasculogenesis of HUVECs in the presence of hMSCs. Therefore, as shown in Figure 3a, without supporting cell migration such as collagen gel and GelMA, or cell-cell interaction, such as in transwells, vasculogenesis of HUVECs cannot be initiated or maintained. Taken together, using vasculature formation by HUVECs and hMSCs in the 3D tissues as an example, we demonstrated that this platform can be utilized to explore the physiology of cell-ECM and cell-cell interactions in the context of drug treatments.

**Figure 5.**
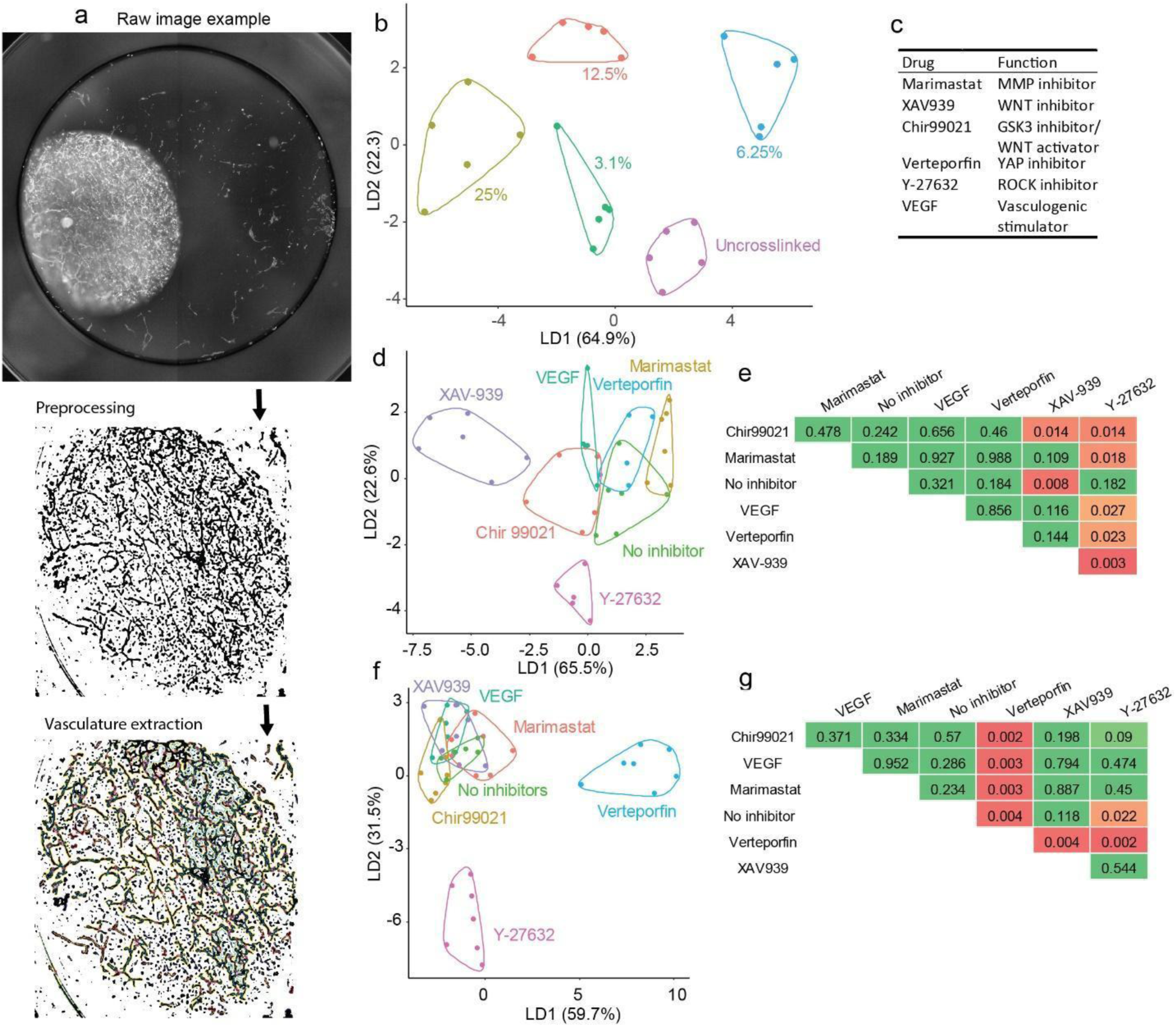
Phenotypic screening of the vasculature formations under different conditions. a, Workflow of whole-mount image processing. Preprocessing was performed using Python with OpenCV. Vasculature multivariable extraction was performed using FIJI (ImageJ) with the Angiogenesis analyzer plugin. b, LDA analysis of vasculature generated within collagen fibers with different crosslinking percentages. c, Drugs used to alter the interactions of cells with different mechanisms to impact blood vessel formation. d, LDA results on Day 3 of vasculature formation with drugs added immediately after tissue generation. e, relative p values pairwised PERMANOVA of d. f, LDA results on Day 6 of vasculature formation with drugs added 3 days after tissue formation. g, relative p values pairwised PERMANOVA of d. Pairwised p values in e and g are color-coded from green to red indicating p > 0.05, 0.01< p < 0.05, and P < 0.01, respectively.

### Screening for autism spectrum disorder using the constructed tissues

We next tested whether this platform can be expanded to other tissue models through phenotypic screening which are traditionally difficult to distinguish. Autism spectrum disorder (ASD) is a complicated neurological disorder that affects patient communication and social behavior. Due to the multiple cell type involvement and complexity of the genes involved and their regulation, traditional homogeneous 2D cell culture models with gene manipulation for ASD research are limited. Therefore, building a platform to improve our understanding of ASD holds significant interest, with major opportunities for basic research, preclinical study, and drug discovery communities. Recent studies found that neural progenitor cells (NPCs) derived from ASD patients may display aberrant cell-cell^24–26^ and cell-ECM^27^ interactions. Therefore, we sought to test if the fast proliferation observed in NPCs generated from ASD donors compared with age/gender-matched control individuals in 2D culture^28^ can be observed in the collagen fiber platform.

We included 7 ASD NPCs (namely, Able, Acai, Ahoy, Apex, Aqua, Arch, Avid), and four non-ASD control NPCs (namely, Chap, Clay, Clue, BJ4).^28^ After plating NPCs containing collagen fiber tissues in the 96-well plate, cells healthily proliferated as seen through LSCM images (Figure 6a). We also quantitatively monitored both the DNA contents and tissue contraction degrees on Day 1, 4 and 7, then normalized all values to those of Day 1 (Figure 6b). DNA content represents NPC proliferation. After 7-day culture, all four NPCs from non-ASD control donors proliferated from 50% to 120% (Figure 6c), whereas NPCs from ASD donors proliferated from 75% to 250% and four of them showed more than 150% proliferation. This result is consistent with previous findings that some NPCs from ASD probands proliferate slightly faster than those from control individuals.^28^ Tissue contraction was measured by residual sizes of tissues in the plate during culture (Figure 6d), which is attributed to the cell-ECM interactions.^29^ Few of the NPCs from control individuals contracted the collagen fibers during a 7-day culture. In contrast, most NPCs from ASD probands perceivably contracted collagen fibers to varying extents. LDA analysis showed clustering of the various groups (Figure 6e). All NPCs from non-ASD control individuals were clustered together. Contrarily, all NPCs from ASD probands scattered on the plot. Excitingly, most NPCs from ASD donors were well separated from the controls except Apex and Able. Understandably, ASD is a spectrum disorder with varied etiology and comorbidities, which may explain the different phenotypical similarities between ASD and controls. We then correlated the LDA scores (LD1-LD4; see Supplementary Table 4 for full correlations of individual variables, LDA scores, and patient information) with the donor information^28^ (Figure 6f). Interestingly, LD1 moderately correlates with the brain volume at years 2-4, and is weakly associated with donor IQ at biopsy. LD2 strongly correlates with the IQ at biopsy and moderately correlates with the brain volume at years 2-4. On the other hand, LD4 exhibited a correlation with the age at biopsy. Previous study revealed the correlation between ASD and WNT signaling, neuron development, synapse morphology/function and organ morphogenesis.^30^ Current development collected detectable signals to reflect these differences and signaled the phenotypic variations of NPCs from ASD probands compared to those from non-ASD donors, showing the promise of using this platform to phenotypically model disorders or diseases with complicated mechanisms *in vitro*.

**Figure 6.**
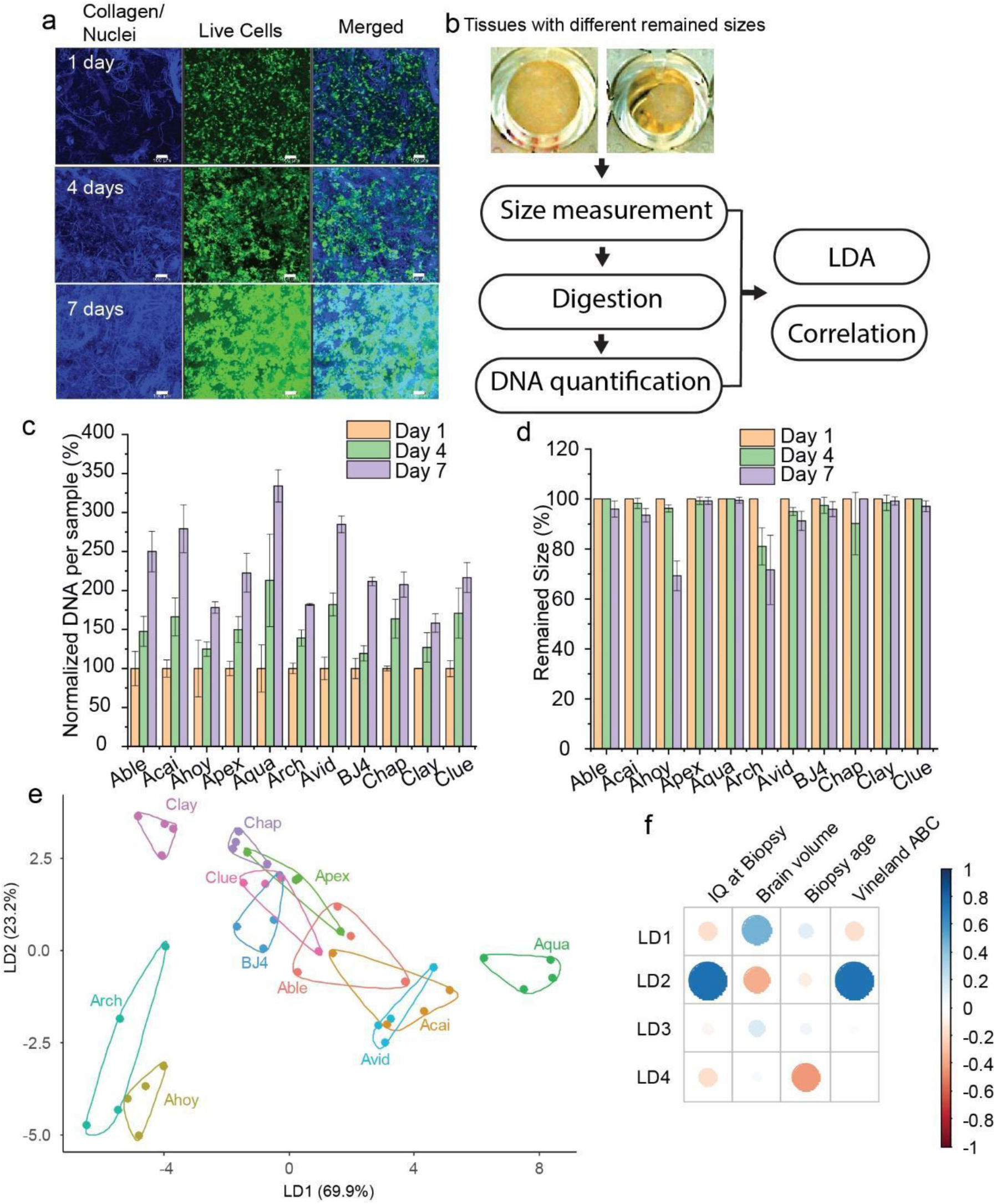
Neural progenitor cells from ASD and control donors grow differently in collagen fiber platforms. a, Z-stack LSCM image of NPC in the collagen fiber construct for 7-day culture. Blue: autofluorescence of collagen fiber and nuclei stained by Hoechst 33342; green: live cells stained by fluorescein diacetate. Bars: 100 μm. b, workflow of the measurements and analysis. c, Normalized DNA content assays for NPCs from ASD and control donors growing in the collagen fiber 3D constructs for 7 days. DNA contents after one day were set to 100%. d, Normalized sizes of 3D constructs cultured for 7 days. The original size was set to 100%. e, Plot of NPCs with their proliferation and tissue contraction using LDA scores LD1 and LD2. f, Correlations of LDA scores (LD1-LD4) and donor IQ at biopsy, brain volume at 2 to 4 years old, age at biopsy, and Vineland adaptive behavior composite (Vineland ABC) at biopsy. The scores are between -1 and 1. The larger the absolute value is, the stronger the correlation is (positively or negatively).

## Discussion

In this work, we adapted the water-lay technique for paper and non-woven fabric manufacturing and established a 3D tissue fabrication platform co-laid with native collagen fibers and cells. The collagen fibers are slurry-like in an aqueous solution and can be easily manipulated using a syringe or pipette and potentially a device with higher throughput. The formation of tissues using collagen fibers is rapid and scalable. A short centrifugation following mixing co-precipitated collagen fibers with cells. After that, cells and collagen fibers assemble producing tissue that is soft but robust. Through various experiments, we have demonstrated that collagen fibers maintain cell-cell and cell-ECM interactions, allowing the distinction of phenotypic changes in cells resulting from different treatment conditions. Of particular interest, with multivariate analysis of 20 variables out of the high-content images, this screening platform detected the impact of the collagen fiber crosslinking and the roles of WNT, YAP, and ROCK pathways in the initiation and formation of vasculogenesis. In addition, when growing NPCs in this platform, the phenotypic differences between ASD and non-ASD donors were distinctively detected, showing the promise of this platform to model disorders or diseases with complicated or unclear mechanisms.

Despite the promising potential, we cannot ignore the limitations of the current prototype. For example, our current preparation strategy did not show detection of other ECM molecules other than collagens and keratin. This component loss is indeed a benefit if xenogeneic tissues are used to prepare the collagen fibers, which can diminish the immunogenicity. If human tissue sources are used to preserve other ECM molecules, such as collagen-bound growth factors, the enzymatic treatment may be minimized, and more physical treatments can be employed. Another potential challenge is to expand this platform to a broader range of cell and tissue types. Right now, the construction conditions of each tissue type, such as the cell amount, media type, collagen usage, centrifugation speed, and culture duration, have to be studied individually. Expansion of the platform to another tissue would be labor-intensive. In addition, although we successfully demonstrated the functionality of *in vitro*-formed vasculature and neural tissues, extensive development and validation of complicated tissues will be required. Despite these concerns, we hope that this rapid and scalable phenotypic screening platform will chart alternative approaches to tissue and disease/disorder modeling, drug discovery, pharmacology/toxicology study, and other applications.

## Author contributions

RT conceptualized the project. EA supervised the project. RT performed the investigations. AD, KU, MR, AD, and CC performed vasculogenesis-related research, CF and AWB performed neural stem cell-related research. LL and SE worked on the material characterization. RT wrote the original draft. All authors edited the draft.

## Acknowledgments

The authors gratefully acknowledge funding from the National Institutes of Health’s National Institute of Arthritis and Musculoskeletal and Skin Diseases (R01AR063194, R01AR074948; EA), National Institute of Biomedical Imaging and Bioengineering (R01EB023907; EA), and National Heart, Lung, and Blood Institute (T32HL134622, T32HL007829; RT), UIC CCTS Pilot Grant (EA and RT), UI Health Diabetes Research Philanthropic Fund (CC and RT).

## Data availability

All data generated or analyzed during this study are included in this published article (and its supplementary information files).

## References

1. Vincent, F., Nueda, A., Lee, J., Schenone, M., Prunotto, M., and Mercola, M. (2022). Phenotypic drug discovery: recent successes, lessons learned and new directions. Nature Reviews Drug Discovery 21, 899–914.

2. Berg, E.L. (2021). The future of phenotypic drug discovery. Cell Chemical Biology 28, 424–430.

3. Langhans, S.A. (2018). Three-Dimensional in Vitro Cell Culture Models in Drug Discovery and Drug Repositioning. Front Pharmacol 9, 6–6.

4. Friese, A., Ursu, A., Hochheimer, A., Schöler, H.R., Waldmann, H., and Bruder, J.M. (2019). The Convergence of Stem Cell Technologies and Phenotypic Drug Discovery. Cell Chemical Biology 26, 1050–1066.

5. Brancato, V., Oliveira, J.M., Correlo, V.M., Reis, R.L., and Kundu, S.C. (2020). Could 3D models of cancer enhance drug screening? Biomaterials 232, 119744.

6. Madl, C.M., LeSavage, B.L., Dewi, R.E., Dinh, C.B., Stowers, R.S., Khariton, M., Lampe, K.J., Nguyen, D., Chaudhuri, O., Enejder, A., and Heilshorn, S.C. (2017). Maintenance of neural progenitor cell stemness in 3D hydrogels requires matrix remodelling. Nature Materials 16, 1233–1242.

7. Zhu, M., Li, W., Dong, X., Yuan, X., Midgley, A.C., Chang, H., Wang, Y., Wang, H., Wang, K., Ma, P.X., et al. (2019). In vivo engineered extracellular matrix scaffolds with instructive niches for oriented tissue regeneration. Nature Communications 10, 4620.

8. Duval, K., Grover, H., Han, L.-H., Mou, Y., Pegoraro, A.F., Fredberg, J., and Chen, Z. (2017). Modeling Physiological Events in 2D vs. 3D Cell Culture. Physiology (Bethesda) 32, 266–277.

9. Fatehullah, A., Tan, S.H., and Barker, N. (2016). Organoids as an in vitro model of human development and disease. Nature Cell Biology 18, 246–254.

10. Roskelley, C.D., Srebrow, A., and Bissell, M.J. (1995). A hierarchy of ECM-mediated signalling regulates tissue-specific gene expression. Current opinion in cell biology 7, 736–747.

11. van Wachem, P.B., and van Luyn, M.J.A. (2001). Collagen Derived Materials. In Encyclopedia of Materials: Science and Technology, K.H.J. Buschow, R.W. Cahn, M.C. Flemings, B. Ilschner, E.J. Kramer, S. Mahajan, and P. Veyssière, eds. (Elsevier), pp. 1313–1314.

12. Sasaki, S., Ikeda, T., Okihara, S.-i., Nishimura, S., Nakadate, R., Saeki, H., Oki, E., Mori, M., Hashizume, M., and Maehara, Y. (2019). Principles and development of collagen-mediated tissue fusion induced by laser irradiation. Scientific Reports 9, 9383.

13. Doillon, C.J., Dunn, M.G., Bender, E., and Silver, F.H. (1985). Collagen Fiber Formation in Repair Tissue: Development of Strength and Toughness. Collagen and Related Research 5, 481–492.

14. Xie, J., Bao, M., Bruekers, S.M.C., and Huck, W.T.S. (2017). Collagen Gels with Different Fibrillar Microarchitectures Elicit Different Cellular Responses. ACS Appl Mater Interfaces 9, 19630–19637.

15. Tang, R., Liao, X.-P., and Shi, B. (2008). Heterogeneous Gold Nanoparticles Stabilized by Collagen and Their Application in Catalytic Reduction of 4-Nitrophenol. Chemistry Letters 37, 834–835.

16. Mills, S.J., Cowin, A.J., and Kaur, P. (2013). Pericytes, mesenchymal stem cells and the wound healing process. Cells 2, 621–634.

17. Caporali, A., Martello, A., Miscianinov, V., Maselli, D., Vono, R., and Spinetti, G. (2017). Contribution of pericyte paracrine regulation of the endothelium to angiogenesis. Pharmacology & Therapeutics 171, 56–64.

18. Chen, Q., and Wang, Y. (2020). The application of three-dimensional cell culture in clinical medicine. Biotechnology Letters 42, 2071–2082.

19. Yap, K.N., and Zhang, Y. (2021). Revisiting the question of nucleated versus enucleated erythrocytes in birds and mammals. American Journal of Physiology-Regulatory, Integrative and Comparative Physiology 321, R547–R557.

20. Carpentier, G., Berndt, S., Ferratge, S., Rasband, W., Cuendet, M., Uzan, G., and Albanese, P. (2020). Angiogenesis Analyzer for ImageJ — A comparative morphometric analysis of “Endothelial Tube Formation Assay” and “Fibrin Bead Assay”. Scientific Reports 10, 11568.

21. Dejana, E., and Kühl, M. (2010). The Role of Wnt Signaling in Physiological and Pathological Angiogenesis. Circulation Research 107, 943–952.

22. Azad, T., Ghahremani, M., and Yang, X. (2019). The Role of YAP and TAZ in Angiogenesis and Vascular Mimicry. Cells 8, 407.

23. Scott, K.E., Fraley, S.I., and Rangamani, P. (2021). A spatial model of YAP/TAZ signaling reveals how stiffness, dimensionality, and shape contribute to emergent outcomes. Proceedings of the National Academy of Sciences 118, e2021571118.

24. Wang, M., Wei, P.-C., Lim, C.K., Gallina, I.S., Marshall, S., Marchetto, M.C., Alt, F.W., and Gage, F.H. (2020). Increased Neural Progenitor Proliferation in a hiPSC Model of Autism Induces Replication Stress-Associated Genome Instability. Cell Stem Cell 26, 221–233.e226.

25. 25. Karpowicz, P., Willaime-Morawek, S., Balenci, L., DeVeale, B., Inoue, T., and van der Kooy, D. (2009). E-Cadherin regulates neural stem cell self-renewal. J Neurosci 29, 3885–3896.

26. Kang, P.H., Kumar, S., and Schaffer, D.V. (2017). Making way for neural stemness. Nature Materials 16, 1174–1176.

27. Lewis, E.M.A., Meganathan, K., Baldridge, D., Gontarz, P., Zhang, B., Bonni, A., Constantino, J.N., and Kroll, K.L. (2019). Cellular and molecular characterization of multiplex autism in human induced pluripotent stem cell-derived neurons. Molecular Autism 10, 51.

28. Marchetto, M.C., Belinson, H., Tian, Y., Freitas, B.C., Fu, C., Vadodaria, K.C., Beltrao-Braga, P.C., Trujillo, C.A., Mendes, A.P.D., Padmanabhan, K., et al. (2017). Altered proliferation and networks in neural cells derived from idiopathic autistic individuals. Molecular Psychiatry 22, 820–835.

29. Liu, A.P., Chaudhuri, O., and Parekh, S.H. (2017). New advances in probing cell–extracellular matrix interactions. Integrative Biology 9, 383–405.

30. Fu, C., Ngo, J., Zhang, S., Lu, L., Miron, A., Schafer, S., Gage, F.H., Jin, F., Schumacher, F.R., and Wynshaw-Boris, A. (2023). Novel correlative analysis identifies multiple genomic variations impacting ASD with macrocephaly. Human Molecular Genetics 32, 1589–1606.

